# Precision of the Integrated Cognitive Assessment for the assessment of neurocognitive performance in athletes

**DOI:** 10.1101/2023.03.22.533746

**Authors:** Daniel J. Glassbrook, Paul L. Chazot, Karen Hind

## Abstract

Choice reaction time tests are commonly used for the assessment of cognitive function, and may be useful to assess the effect of sport participation. This study investigated the precision of the Integrated Cognitive Assessment (ICA; Cognetivity Neurosciences Ltd., Vancouver, Canada) test for the assessment of cognitive function in athletes. Thirty-one participants volunteered to take part in this study, from both contact (*n* = 22) and non-contact sports (*n* = 9). Participants performed the ICA test consecutively both before and after normal training session to simulate resting and post-sport conditions. Precision errors, relationships (Pearson’s r), and internal consistency (Cronbach’s Alpha) were calculated for three variables, ICA Index (overall information processing ability), ICA Speed (information processing speed) and ICA Accuracy (information processing accuracy). ICA precision errors [root mean squared-standard deviation, RMS-SD (coefficient of variation, %CV)] pre-sport were ICA Index: 5.18 (7.14%), ICA Speed: 3.98 (4.64%), and ICA Accuracy: 3.64 (5.00%); and post-sport were ICA Index: 3.96 (4.94%), ICA Speed: 2.14 (2.32%), and ICA Accuracy 3.40 (4.25%). The ICA test demonstrates high in-vivo precision with all variables except ICA Index (7.14%) demonstrating an acceptable precision error of ≤5% %CV. All variables demonstrated strong relationships between consecutive tests pre- and post-sport (r ≥ 0.8) except for the ICA Index post-sport which demonstrated a moderate (r ≥ 0.5) relationship. The ICA Index demonstrated good internal consistency (α ≥ 0.8) for both pre-and post-sport. The ICA Speed and ICA Accuracy variables demonstrated excellent internal consistency (α ≥ 0.9) for both pre-and post-sport. The ICA test is suitable for the assessment of cognitive function pre- and post-sport.

## 1.0 Introduction

The brain plays a vital role in sports participation, and is responsible for decision making, and good decision making is key to success in sport [1]. Athletes are required to demonstrate high levels of technical and perceptual decision making ability, all while under pressure and fatigue [2]. Assessing cognitive function, such as information processing and decision making ability is a valuable tool which may quantify psychological ability in athletes. Additionally, assessing cognitive function may give insight into the effect of sport participation on the brain, for example through the effect on reaction time and decision making ability. There are many positive effects of sport and exercise participation, including cognitive benefits such as neuroplasticity, the process of adaptive structural and functional changes to the brain [3]. However, injury is an innate part of sport participation due to the physical nature of sport, and head impacts which may impair cognitive function are associated with participation in contact sports.

One proposed method to assess the effects of possible deleterious effects of head impacts sustained during normal sport participation, and cognitive function clinically is via information processing and reaction time tests. Reduced information processing ability, including memory, is linked with head impacts [4-6]. Indeed, altered mental state and confusion are also symptoms associated with head impacts that can negatively affect memory and information processing ability [7]. Additionally, information processing speed underpins several conditions of cognitive dysfunction, for example, multiple sclerosis [8, 9] and Alzheimer’s disease [10].

Slower reaction time is a commonly used indicator of cognitive change following head impacts [11, 12]. Two common types of reaction test include measurement of simple reaction time (SRT) or choice reaction time (CRT) [13]. SRT is recorded when there is only one possible stimulus (signal) and one possible response (action), for example tapping anywhere on a screen when any image appears. In CRT tasks there are two or more possible stimuli, each of which requires a quite different response, for example, tapping on the left of the screen when an image of an object appears on the screen, and tapping on the right when an image of an animal appears on the screen. SRT tests such as a weighted object drop and catch [14], ruler drop test [15], and somatosensory assessment, where the participant held a device similar to a computer mouse and reacted to a vibration applied to one or both the index or middle finger, and responded by clicking with the finger (or both) that received the vibration [16], have all been used to show slower reaction time in participants that have experienced head impacts. Computerised tests have also been shown to be sensitive and able to determine athletes who experienced head impacts versus those without, for example the CogSport choice reaction time test and Immediate Post-Concussion Assessment and Cognitive Testing (imPACT) [17-20]. The CogSport choice test takes approximately 20 minutes, and consists of seven tasks including a SRT, CRT and complex reaction time task, as well as tests of visual and working memory, all designed as card games [21]. The imPACT is performed with a computer screen and mouse and typically takes 20-25 minutes and comprises of six test modules assessing attention, memory, processing speed and reaction time, through tests such as word memory and symbol matching [22]. Interestingly, reaction time has also been used to show the lasting effect of head impacts on reaction time, with reaction time scores only returning to baseline at 10 to 14 days post injury [15], and up to 21 days post injury [23].

The Integrated Cognitive Assessment (ICA; Cognetivity Neurosciences Ltd., Vancouver, Canada) [24, 25], is a newly developed method for the assessment of cognitive function, and may be applicable to the assessment of cognitive function in athletic populations. The ICA is a short computerised cognitive test of cognitive function via an assessment of information processing (CRT) speed based on a rapid categorisation task [24, 26]. The ICA test can be completed in a short amount of time (∼5 minutes), on a handheld device such as an iPhone or iPad, and is therefore able to be administered easily pre- and post-sport. Performing an assessment of cognitive function such as the ICA pre-sport in a rested state effectively provides a baseline measure of cognitive function. Performing a test of cognitive function post-sport then can be compared to the baseline measure of pre-sport and determine the effect of sport participation on cognitive function. Additionally, the ICA has been shown to accurately detect mild cognitive impairment and be moderately to highly correlated with other popular pen-and-paper cognitive tests such as the Montreal Cognitive Assessment (Pearsons r = 0.58) and Addenbrooke’s Cognitive Examination (Pearsons r = 0.62) cognitive tests [24].

The ICA has been shown to accurately measure cognitive impairment in patients in the early stages of dementia [24]. However, to date, no known study has investigated the intra-day precision of the ICA test. Therefore, the purpose of this study was to determine the same-day, in-vivo (i.e., within whole living organisms; within people) precision of the ICA test to assess cognitive function. It is hypothesised that the ICA will demonstrate acceptable precision, and suitability for the assessment of cognitive function pre- and post-sport.

## 2.0 Materials and Methods

### 2.1 Participants

Thirty-one participants volunteered to take part in this study. The minimum number of participants required for precision analysis with two consecutive tests is thirty [27]. Participant characteristics are presented in Table 1. Participants were recruited from March to May 2022, from the University sporting facility via advertisement flyer, and from the wider sporting community through existing relationships with sporting clubs. After initially expressing interest to take part in the study, participants were provided with an information sheet and given at least 24 hours to consider their participation in the study before a follow up was sent by the lead researcher. After confirmation of eligibility, participants were then booked in at a time and date of their convenience for data collection. Participants were eligible for participation if they were a current contact sport or non-contact sport athlete (Table 2) aged 18-40 years, and healthy; having no underlying medical issues that affect participation in sport. Participants were excluded if they were injured, pregnant, or suffering from post-concussion syndrome. Post-concussion syndrome is defined as the collective experience of symptoms after sustaining a concussion, such as headaches, fatigue, depression, anxiety and irritability [28, 29]. This study was approved by the Durham University Sport and Exercise Sciences Ethics Committee (reference: SPORT-2022-01-07T10_44_59-srhd22;), and written informed consent was provided by each participant prior to participation. Participant characteristics, demographics and athletic history were collected via standardised questionnaire directly after consent to participate was obtained.

**Table 1:**
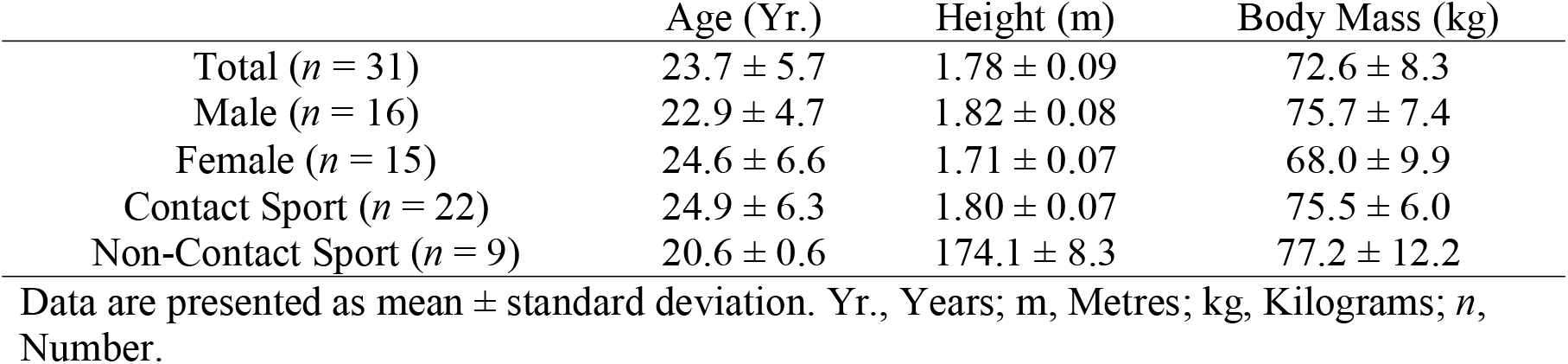
Participant Characteristics

**Table 2:** Sports Breakdown

| Contact Sports ($n = 22$) | | Non-Contact Sports ($n = 9$) | |
| --- | --- | --- | --- |
| Rugby Union ($n = 7$) | Semi-professional,<br>Amateur | Touch Rugby<br>($n = 5$) | Amateur |
| Boxing ($n = 6$) | Amateur | Athletics ($n = 4$) | Amateur |
| Muay Thai (Kickboxing) ($n = 5$) | Professional, Amateur | | |
| Indoor Football ($n = 4$) | Amateur | | |

$n$ , Number.

### 2.2 Procedures

To simulate resting- and post-sport conditions, participants performed the ICA test (version 1.6.0 or 1.7.0) before and after a normal training session for their respective sports. Data collection was performed in a quiet room to minimise distractions, with a maximum of two participants (separated within the room) performing the test at a time. Prior to their sports training, and within an hour of the training start time, participants completed two consecutive ICA tests. The participants then completed a normal training session and then two consecutive ICA tests again, within an hour of the training finishing.

### 2.3 The Integrated Cognitive Assessment (ICA)

The ICA has been described in detail in published works [26, 30]. In summary, the ICA is a short computerised cognitive test of cognitive function via an assessment of information processing (CRT) speed based on a rapid categorisation task, and in contrast to the CogSport and imPACT tests is independent of language or colour. The ICA in this study was completed on an iPad, with each test taking approximately five minutes, considerably shorter than the 20+ minute CogSport and imPACT tests; the imPACT test of which is also not portable (completed on a computer with screen and mouse). Each ICA test utilises the brains powerful reaction to animal stimulus [31] and comprises of 100 natural images, with 10 additional practice images provided in a practice section prior to the test. The practice images are not included in the test score. The images are a mixture of animals (e.g., birds, fish, mammals), or non-animals (e.g., objects, food, vehicles), and are presented in rapid succession. Images appear for 100ms, followed by a 20ms inter-stimulus interval and a 250ms dynamic noisy mask. Participants react to the image by tapping with their thumb on the screen; on the right-hand side to select ‘animal’ and on the left-hand side of the screen to select ‘non-animal’. Participants were instructed to perform the test as quick as possible, and these instructions were given both verbally and presented on the screen. There are no associated risks with completing the ICA, and the test provides three variables: ICA Speed, the response reaction times in trials they responded correctly; ICA Accuracy, choice accuracy, the percentage of correct test responses; and ICA Index, overall information processing ability, a combination of ICA Speed and ICA Accuracy. Each variable is calculated by the following equations [24, 30]:

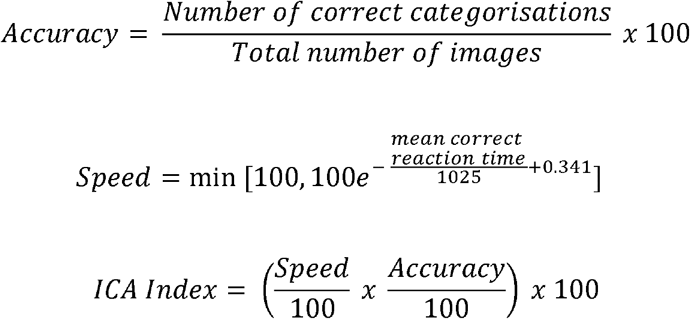

There is a speed-accuracy trade-off in reaction test performance, and often scoring higher in either speed or accuracy is achieved at the expense of the other capacity [32, 33]. To combat the potential negative reflection on overall information processing ability from a poor speed or accuracy score, a common solution is the inverse efficiency score [34], whereby speed and accuracy are combined into a single score. In the case of the ICA, this concept is applied and manifests as the ICA Index variable. Test results are recorded via report immediately after test completion.

ICA Accuracy, ICA Speed and ICA Index are represented by Arbitrary Units and results of each equation are transformed and scored by a standardised scale ranging from 0 to 100. I.e., a Speed score closer to 100 indicates faster reaction times, while a Speed score closer to 0 indicates slower reaction times; an Accuracy score closer to 100 indicates higher accuracy, and an Accuracy score closer to 0 indicates lower accuracy; and an ICA Index score closer to 100 represents greater overall information processing ability, whereas an ICA Index score closer to 0 indicates poorer overall information processing ability. Additionally, the specific components included in the Speed equation (i.e., division by 1025 and addition of 0.341) further serve to standardise and scale the Speed score. These additional components are determined empirically to ensure that the Speed scores fall within the desired range of 0 to 100 for the majority of participants. Natural Log transformation further aids in normalizing the distribution of scores.

### 2.4 Statistical Analysis

All data analysis was performed in Microsoft Excel (2016). Raw anonymised data for ICA Index, ICA Speed, and ICA Accuracy were extracted and exported to Microsoft Excel for analysis [35]. Throughout the study the lead author had access to information that could identify individual participants, however, all analysis was performed on anonymised data. Precision has been previously used to define test suitability in similar cognitive tests to the ICA, such as the CogSport choice reaction time test [36]. Precision of ICA scores and least significant change (LSC) were calculated at the 95% confidence level. Precision was determined as root mean square standard deviation (RMS-SD), coefficient of variation (CV), and percentage CV (%CV). RMS-SD represents the sample standard deviation of the differences between predicted values and observed values, and is calculated via the following formulae, where SD represents standard deviation and *n* represents the number of participants:

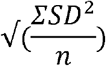

The %CV expresses test variation relative to the mean of two tests and is corrected for small sample bias, and was defined as acceptable <5% [37]. The LSC represents a true meaningful change was calculated from the precision errors (LSC = RMS-SD ^*^ 2.77). Pearson’s correlation coefficient was calculated for each variable to determine reliability and the strength of the relationship between consecutive tests, with 0.2, 0.5, and 0.8 representing weak, moderate, and strong correlations, respectively [38]. Cronbach’s Alpha was calculated for each variable to determine internal consistency, with 0.9, 0.8, 0.7, 0.6, 0.5 and < 0.5 representing excellent, good, acceptable, questionable, poor and unacceptable internal consistency, respectively [39].

## 3.0 Results

Results of the precision analysis for each ICA variable pre- and post-sport are presented in Table 3. All variables except for ICA Index pre-sport had a precision error of ≤5% %CV. LSC results are presented in Table 4. Pearson correlation coefficient data is presented in **Error! Reference source not found**.. All variables demonstrated strong relationships between consecutive tests pre- and post-sport, except for the ICA Index post-sport which demonstrated a moderate relationship. Cronbach’s Alpha data is presented in Table 5. The ICA Index demonstrated good internal consistency for both pre-and post-sport. The Speed and Accuracy variables demonstrated excellent internal consistency for both pre-and post-sport.

**Table 3:** Precision Analysis Results

| Variable | Precision ($n = 31$) | | |
| --- | --- | --- | --- |
|  | RMS-SD | CV | %CV |
| *Pre* |  |  |  |
| ICA Index | 5.18 | 0.07 | 7.14 |
| ICA Speed | 3.98 | 0.05 | 4.64 |
| ICA Accuracy | 3.64 | 0.05 | 5.00 |
| *Post* |  |  |  |
| ICA Index | 3.96 | 0.05 | 4.94 |
| ICA Speed | 2.14 | 0.02 | 2.32 |
| ICA Accuracy | 3.40 | 0.04 | 4.25 |

n, Number; ICA, Integrated Cognitive Assessment; RMS-SD, Root Mean Square Standard Deviation; CV, Coefficient of Variation; %, Percentage.

**Table 4:** Least Significant Change Results

| Variable | LSC ($n = 31$) | | |
| --- | --- | --- | --- |
|  | RMS-SD | CV | %CV |
| *Pre* |  |  |  |
| ICA Index | 14.36 | 0.20 | 19.78 |
| ICA Speed | 11.01 | 0.13 | 12.86 |
| ICA Accuracy | 10.09 | 0.14 | 13.9 |
| *Post* |  |  |  |
| ICA Index | 10.96 | 0.14 | 13.7 |
| ICA Speed | 5.94 | 0.06 | 6.43 |
| ICA Accuracy | 9.43 | 0.12 | 11.78 |

LSC, Least Significant Change; n, Number; ICA, Integrated Cognitive Assessment; RMS-SD, Root Mean Square Standard Deviation; CV, Coefficient of Variation; %, Percentage.

**Table 5:** Relationships between consecutive tests

| Variable | Test 1 | Test 2 | Correlation (r) | Cronbach's Alpha ($\alpha$) |
| --- | --- | --- | --- | --- |
| *Pre* |  |  |  |  |
| ICA Index | 76.6 ± 8.3 | 82.2 ± 6.1 | 0.82 | 0.88 |
| ICA Speed | 88.6 ± 8.1 | 92.3 ± 7.3 | 0.85 | 0.91 |
| ICA Accuracy | 86.9 ± 8.9 | 89.2 ± 6.2 | 0.87 | 0.90 |
| *Post* |  |  |  |  |
| ICA Index | 81.5 ± 7.0 | 83.0 ± 6.7 | 0.68 | 0.81 |
| ICA Speed | 94.8 ± 5.2 | 95.6 ± 4.6 | 0.82 | 0.90 |
| ICA Accuracy | 86.1 ± 9.1 | 87.0 ± 8.3 | 0.85 | 0.92 |

n, Number; ICA, Integrated Cognitive Assessment; Data are presented as mean ± standard deviation.

## 4.0 Discussion

The purpose of this study was to determine the same-day, in-vivo precision of the ICA test to assess cognitive function. The results of this study support the ICA as a tool with acceptable precision to measure changes in cognitive ability pre- and post-sport, confirming the hypothesis. All ICA variables in this study, except for ICA Index pre-sport demonstrated a precision error of ≤5% %CV. All variables demonstrated moderate to strong relationships and good to excellent internal consistency between consecutive tests pre- and post-sport. The ICA may be clinically relevant for the assessment of cognitive ability, and the effect of sport on cognitive ability.

The higher ICA Index precision score (7.14 %CV) pre-sport than post-sport (4.94 %CV) in this study may be explained by the larger average difference between test one and test two pre-sport (76.6 ± 8.3 to 82.2 ± 6.1), compared to a smaller average difference in ICA Index scores between test one and test two post-sport (81.5 ± 7.0 to 83.0 ± 6.7). This is further exemplified by a larger ICA Index RMS-SD pre-sport than post-sport (5.18 and 3.96, respectively), which indicates more variance in observed data around the mean. This result may be due to an increased level of comfort with the test from the first pre-sport ICA test to the subsequent test, and possibly a learning effect. However, this is in contrast to previous work which showed no learning effect for the ICA test in healthy participants and those diagnosed with dementia [24]. Additionally, a strong relationship (r = 0.82) and good internal consistency (α = 0.88) between ICA tests and the ICA Index variable was observed pre-sport (Test 1 and Test 2; Table 5), further suggesting a minimal learning effect, and supporting the test as reliable.

All variables showed greater precision post-sport compared to pre-sport. This may be due to the many positive physiological benefits that exercise has, such as an increase in blood flow to muscles and brain [40], structural and functional changes in the brain [3], and increases information processing ability [41]. Indeed, improvements in cognitive function after a bout of exercise is supported by previous research [42, 43].

Previous research looking at precision in a similar cognitive test to the ICA, the CogSport choice reaction time test, has shown lower %CV for mean choice reaction time (speed) (1.4 %CV), and higher %CV for choice reaction time accuracy (11.4 %CV) [36]. These results are interesting as the ICA is shown to be less precise in measuring reaction speed (2.32 - 4.64 %CV vs 1.4 %CV), however, the ICA test is shown to be more precise in terms of accuracy (4.25 – 5.00 %CV vs 11.4 %CV). These results may indicate that the test you adopt needs to be specific to the variable of interest (i.e., speed or accuracy), however, this should be negated in the case of the ICA via the ICA Index variable as an inverse efficiency score [34], whereby speed and accuracy are combined into a single score. The contrasting results between the present study and that of Straume-Naesheim, Andersen and Bahr [36] may be due to the populations used; the CogSport test was used in elite football players only, whereas only a small percentage of the participants in the present study are practicing professionally (table 2). The present study recruited participants from a variety of sports, each with their own decision making and reaction characteristics, in comparison to only football. Additionally, a greater number of participants (*n* = 289) performed the CogSport test in comparison to the ICA in the present study (*n* = 31). However, the administration procedure of the CogSport test was similar to the administration of the ICA test, and was also administered under controlled conditions, i.e., in a quiet room to minimise distractions and with more than one participant performing the test at a time.

A limitation to this study is that information about the specific duration an intensity of training sessions undergone by participants was not recorded. Only that it was a typical or normal training session. i.e., not a session of just running or conditioning, and included practice of the sport itself. Additionally, the optimal intensity and duration of exercise that may impact cognitive function was not assessed. A relatively small sample size was used, although the sample size exceeds the minimum of 30 participants required for precision analysis with two consecutive tests [27], a greater number of participants could strengthen the statistical analysis. Future studies should look to recruit larger numbers, and complete comparisons between contact and non-contact sports, and between amateur and professional level participants.

In conclusion, the ICA is a practical test which can be used to measure cognitive function before and after sport participation. The results of this study support the ICA as a precise (as determined by %CV values) measure of information processing speed and information processing accuracy, and overall information processing ability (ICA Index). The ICA can be used for the assessment of cognitive function, and may be useful as a method to assess the effects of sport on cognitive function.

## Notes

**Conflicts of Interest/Competing Interests:** The authors have no relevant financial or non- financial interests to disclose.

https://figshare.com/articles/dataset/ICA_Precision_Raw_Data_xlsx/22211644

### Competing Interest Statement

The authors have declared no competing interest.

### Summary of Updates

This version of the manuscript has been revised to update the introduction, and referencing style.

